# Cleaning clinical genomic data: Simple identification and removal of recurrently miscalled variants in single genomes

**DOI:** 10.1101/237107

**Authors:** Matthew A. Field, Gaetan Burgio, Jalila Al Shekaili, Simon J. Foote, Matthew C. Cook, T. Daniel Andrews

## Abstract

Identification of sequence variation from short-read sequence data is subject to common-yet-intermittent miscalling that occurs in a sequence intrinsic manner. We identify that recurrent false positive single nucleotide variants are strongly present in databases of human sequence variation and demonstrate how each individual sample generates a unique set of recurrent false positive variants. These recurrent miscalls result from known difficulties aligning short-read sequence data between redundant genomic regions. We could replicate, catalogue and remove three quarters of these recurrent miscalls for any given exome with as little as ten rounds of read resampling, realignment and recalling. The removal of such misleading variants reduces the search space for identification of disease causing variants.

**List of Abbreviations:**

SNV: single nucleotide variant
RFP: recurrent false positive
ENU: N-ethyl-N-nitrosourea

## Background

We have recently seen the misinterpretation of spurious single nucleotide variant (SNV) calls from resequencing of the mouse genome following introduction of targeted mutations using CRISPR-Cas9 technology (1, 2). This has been the subject of interesting debate regarding the reproducibility of results from massively parallel sequencing. Equally as relevant to this discussion is the high clinical value of rare variants to the interpretation of human personal genomes. Genomic sequence can be easily obtained in clinical care but subsequently encounters the difficulty of pathogenic variant identification (3). Generally, pathogenic variants are not found at high population frequencies and rare variants are prioritised in searching for causal mutations (3–5). In this context, trace levels of spurious SNV miscalls from short-read sequencing have a disproportionately large impact (6) and may lead to an incorrect diagnosis of pathogenicity.

The history of mammalian genome sequencing has been a progression towards shorter sequencing reads and now relies heavily on aligning these to a reference genome (7). Highly similar genomic regions are difficult to resolve with this information and read misalignment is a prevailing source of variant miscalls (8). Algorithmically, for example, the Burrows-Wheeler Transformation method implemented by the BWA tool (9) must resort to random read assignment between highly similar regions should their mapping quality score fail to differentiate them.

When mapping short-read data to a reference genome, read misalignment has been identified as the predominant source of incorrect variant calls (8). Misalignment of reads in redundant genomic regions is often highly specific to the given genome sequence from which it is derived. It has remained difficult to appraise the quality of single nucleotide variant sets identified for any given individual genome sequence obtained from short-read sequencing data. A small number of gold-standard reference variant sets indicate that read misalignment often leads to variant miscalls in regions of genomic redundancy.

Identification of sequence variation from short-read sequence data is subject to common-yet-intermittent miscalling that occurs in a sequence intrinsic manner. We could replicate and catalogue these recurrent miscalls for any given exome by only ten rounds of read resampling, realignment and recalling. We identify that recurrent false positives are strongly present in databases of human sequence variation and demonstrate how each individual sample generates a unique set of recurrent false positive variants.

## Data and Methods

### Mouse recurrent false positive variants

Exomes from C57BL6 mice were obtained and variation identified as previously described (10). The frequency of variants was ascertained through counting variant occurrence in a population of 2114 exomes from distinct mice. Recurrent false positive variants were identified as those that were present at frequencies greater than 5% but less that 95%.

### Resampling, realignment and recalling

Random reads were sampled from an input genome sequence in Fasta format with the tool WGSim (WGSim GitHub repository: http://github.com/lh3/wgsim). Synthetic exome sequences for both the mouse genome (mm10) and the human genome (hg37d5) were derived from EnsEMBL BioMart (www.ensembl.org) and consisted of the sequences of all exons and 325bp of padding upstream and downstream of each exon. From these derived sequences, mock sequencing datasets were simulated to contain 80 million, 120bp paired-end reads with random sequencing errors at a frequency of 1%. Sequence datasets were aligned to the chosen reference genome with BWA mem (9). Single nucleotide variants were called with SAMtools (17) (http://www.htslib.org) and GATK (18) (http://gatk.broadinstitute.org) using best-practice methodology and parameter sets.

Individual exome and whole genome sequences were derived as alternative reference sequences using the FastaAlternateReferenceMaker tool from the GATK suite.

### Non-reference mouse and human sequence data

Exomes of the FVB, C3H and CBA mouse strains were derived by the resampling, alignment and recalling procedure from these strain sequences (12, 13).

The genome sequence of the human NA12878 individual was obtained from the Illumina Platinum Genomes resource (<u>ftp://ftp.1000genomes.ebi.ac.uk</u>). Omani exome sequences were obtained with approved hospital consent for the genetic analysis of these individuals and sequenced as described previously (19). The Indigenous Australian whole genome sequences were obtained with appropriate consent granted to SJ Foote (The Australian National University 2014/663).

## Results

### Recurrent false positive variants identified from inbred mice

We have previously generated a large dataset of mouse exomes from inbred C57BL6 mice harbouring random, N-ethyl-N-nitrosourea (ENU) induced mutations (10). In addition to the 30-60 induced mutations present per pedigree, we observed a category of variant calls that recurred at seemingly random sites in an intermittent mode. These did not validate with genotyping and were not heritable (10). We refer to these as Recurrent False Positive (RFP) variants. From 2114 sequenced mouse exomes, we identified at total of 104,303 unique SNV sites, the bulk of which are strain-specific variation, but also include ENU-induced mutations (https://databases.apf.edu.au/mutations/) and RFP variants. Figure 1a shows the frequency distribution of all SNVs identified in this population of exome sequences. Strain-specific variation occurs at a frequency approaching 100% and, conversely, ENU induced mutations were pedigree-specific at very low frequencies (<1%). RFP variants are comparatively fewer and occur at intermediate frequencies between these extremes, conservatively between 5 and 95% in our mouse exome population.

**Figure 1.**
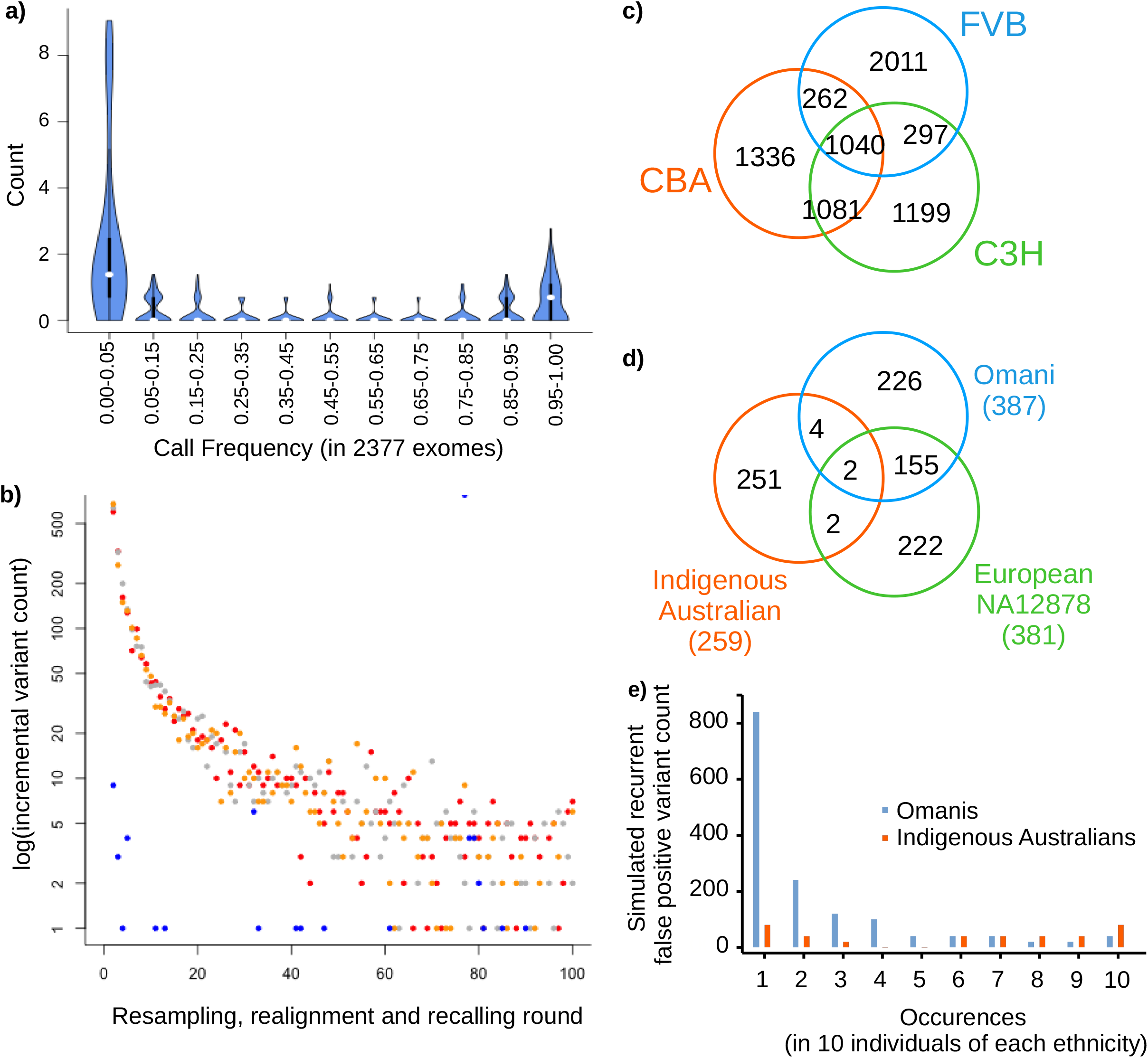
Recurrent false positive variant calls result from difficulties encountered with alignment of short reads to complex mammalian genomes. **a**) The frequencies of observed variant calls from 2314 exomes of inbred mice show an intermediate category of recurrent-yet-intermittent SNVs, between the frequency extremes of fixed strain-specific variation and the rare, pedigree-specific induced mutation. **b**) Recurrent false positive variants can be replicated for a given individual sequence through randomly sampling short reads, realigning these to a reference genome and recalling sequence variants. Variants that are only intermittently called accumulate after multiple cycles of this process and increase in number following an approximately Poisson distribution. Blue dots show the smaller number of recurrent false positive variants obtained from sampling a C57BL6 mouse and realigning it to itself. Greater numbers are obtained through simulation with three non-reference mouse strains FVB (orange), CBA (red), and C3H (grey). **c**) Recurrent false positive variants arise in a sequence-specific manner in three mouse strains. Similarities between these variant sets closely mirror the sequence similarity and relatedness between individual genomes. **d**) Likewise, with human sequences, while a small set of recurrent false positive variants are common between all four individuals, the majority are highly ethnically-dependent to individual-specific. **e**) Further to this, simulation of recurrent false positive variants from a closely related group of Omani individuals indicates that most are individual-specific, with a smaller number being population-specific.

Using these population frequency thresholds, we identified 649,984 variants at 708 unique sites at which RFP variants occur in our data. A distinguishing feature of these variants (compared to randomly chosen single nucleotide positions) is they occur in genomic regions with significantly lower alignability ((11), t-test p<2.2e-16)). Previous work has demonstrated that redundant genomic sequences correlate with low alignability scores and represent one cause of read misalignment (8). Another cause of low alignability is the quality of the reference genome. With variant calling from mouse exomes, we routinely needed to filter 42.5% fewer RFP SNVs when aligning to the improved mm10 reference genome compared to the mm9 reference. Collectively, these factors provide a cogent explanation for the miscalling of RFP variants.

We were able to replicate RFP variant calls from any given single mouse exome, through multiple rounds of resampling of short reads, realigning these with the mouse reference genome and recalling sequence variants. In this process, reads were redundantly and randomly sampled from a full exome sequence to extract 80 million 120bp reads. Each sampled read included randomly introduced sequence errors at a base frequency of 1% to mimic the observed error rate encountered during resequencing with current short-read sequencing technology. Each simulated round from the same exome produced a number of intermittent variant calls that were not reproduced in every round, similar to the RFP variants. Variants called with >95% frequency were predominantly true strain variation and were discarded. Further rounds of this procedure with the same exome incrementally increased the unique RFP variants identified (Figure 1b). The RFP variants obtained in this manner incrementally replicates a subset of the false positive variants observed from actual resequencing. 380 iterations of this resampling, realignment and recalling of the colony C57BL6 genome produced 656 RFP variants, matching 367 (51.8%) of the set observed from actual exome data. A greater number of RFP variants are called when aligning a non-reference mouse strain with the reference. We repeated the resampling, realignment and recalling procedure with three more distant strains FVB (12), CBA and C3H (13) (Figure 1b). The incremental increase of RFP variants with repeated iteration approximated a Poisson distribution with λ = ~1. Just 10 rounds of resampling, realignment and recalling produced 70% of the final total of intermittent variants observed with 100 rounds.

The FVB, CBA and C3H genetic backgrounds produced thousands of RFP variants per genome and these variants variably overlap by between one to two thirds in each mouse strain (Figure 1c). The greater overlap of RFP variants between the more closely related C3H and CBA strains demonstrates how sequence divergence of non-reference strains gives rise to strain-specific variation. The degree of this sequence difference between reference and individual genome substantially contributes to the quantity and distribution of RFP variants in any individual. This has large implications for detecting causal variants in human disease, as the genetic background of any individual will generate a sample-specific set of RFP variants not relevant to their disease.

### Recurrent false positive variants from human sequence data

RFP variants in human sequences could also be identified with the same sampling, realignment and calling method applied to mouse sequences. We generated a catalogue of RFP variants for the HapMap individual NA12878 and compared these with the Genome Aggregation Database (GnomAD; gnomad.broadinstitute.org). Almost, but not all, RFP variants for this individual (96.7%)were present in this database, noting that the NA12878 individual is already present in GnomAD and that RFP variants arise stochastically due to the random sampling of short reads and sequencing error. Further RFP variant cataloguing analysis was also performed on three additional human exomes of ethnicities not dominant in the GnomAD set (an Omani and an Indigenous Australian; Figure 1d). The number of RFP variants in each of these individuals was broadly similar, with an average of 334.5 per individual. The majority are unique to each individual and the proportion of RFP variants that are unique to a given individual varies substantially, though not predictably with, say, sequence divergence from the human reference sequence. The RFP variants per individual were also almost entirely present in the GnomAD database, with population frequencies ranging from common to rare – and importantly, each individual possessed a small number that were unique (Omani, 1.8%; Indigenous Australian, 3.1%; NA12878, 3.3%).

While read mismapping is related to the difficulty an aligner encounters with short-read data, variant callers may differ in their propensity to make RFP variant calls from the same alignment. Our mouse variant calls made with real exomes derived from a mutagenised population of thousands of laboratory mice were performed over time using SAMtools, and we have not replicated these. However, the variant calls for the resampled human data here were made with both SAMtools and GATK (see Methods). Direct comparison was made by making variant calls on the NA12878 individual using the same alignments generated by 30 rounds of read resampling and realignment. Hence, variant callers were working from the same set of read misalignments in each replicate. Interestingly, both variant callers produce RFP variant calls of similar propensity (SAMtools: 334; GATK: 398) but these only overlap by 64 variants, less than a fifth in both cases. Clearly, more four-fifths of RFP variants could be identified and removed by excluding variants not identified by both callers.

The sequence-specificity and covariation with ethnicity of RFP variants was further investigated within single ethnic groups. We repeated the analysis with exomes from ten Omani individuals from pedigrees with an inherited predisposition to autoimmune disease and ten Indigenous Australian individuals with a predisposition to kidney disease. The RFP variants from these individuals of the same ethnicity show more similarity than with other groups (Figure 1e). This is especially apparent for the Indigenous Australian individuals, for whom the RFP variants they hold do not intersect overly with, say, the Omani or NA12878 individuals. The Omani individuals show a strong tendency towards unique RFPs in every individual and less overlap between individuals. Hence there are clear differences between the representative ethnic groups shown here - and this reflects the sequence similarity between the individuals included in each group. In this instance, the catalogue of RFP variants for the Omani individuals is of substantial practical clinical value. Autoimmunity in these individuals could plausibly have been ascribed to predicted-damaging RFP variants in genes with strong associations to lupus (IRF5, LILRB3) or autoimmune hepatitis (C4A). The IRF5 variant in particular was a strong candidate, yet was proven to be miscalled on subsequent genotyping.

## Discussion

Our results identify a class of false variant calls that are an inherent factor in reliably realigning short read sequence information to a complex mammalian genome. This class of variants was shown to arise from analysis of both mouse and human sequences. Significantly, we find that human recurrent false positive SNVs are strongly represented in human population sequence databases, such as GnomAD. Further, these recurrent false positive SNVs may be identified for any given genome sequence through repeated sampling and realignment against a reference sequence. Hence, from this work we show that it is possible to computationally remove the bulk of these spurious SNV calls.

At the heart of these variant miscalls is misalignment of reads between redundant regions of the genome. These redundant regions differ very slightly, so as that the low level of sequencing error inherent in short read data will similar to the true variation that exists between near duplicate sequences in the genome reference. Hence, reads may be stochastically misassigned between these redundant regions and when calling sequence variation, the true differences present in the misaligned reads become called as sequence variation between the re-sequenced and reference genomes. Hence, these miscalls will be recurrent to specific nucleotide sites in any given resequenced genome, yet will recur in an intermittent manner due to the stochastic way in which sequencing error and misalignment occurs. Importantly, this will produce some miscalled variants with what will appear as rare sequence variation when these variants are aggregated in human population variation databases. This mechanism of RFP genesis suggests that their removal can be achieved by both exhaustive cataloguing of RFP variants and/or identifying for a given genome the near-identical regions and the minor sequence differences that exist between them. In practice, the exhaustive cataloguing through similation is simple to achieve. Even more simply, a quick workaround that will remove more than 80% of RFP variants is to remove variant calls that are not replicated by both of SAMtools and GATK. This work-around is a methodology that has increasingly been gaining acceptance for many other purposes also (14) and is further reason for clinical variants to be made from the union of calls made with multiple callers (15, 16).

Simulation and cataloguing of RFP variants is technically simple, but does substantially increase the computation required to perform this analysis for any given genome. Each simulation round requires computation to simulate a population of reads from a given genome, subsequent realignment of these reads and the calling of sequence variation between this alignment and the reference genome. This is effectively (at least) a ten-times increase in computation for a single genome. However, the cost of computing a single exome or genome (in the order of tens of dollars) is relatively small compared to the cost of generating this sequence data (two orders of magnitude more, at current costs). Hence, increasing this compute cost by a factor of ten is inconsequential should this improve the quality of the derived information substantially. Furthermore, the costs of a misidentified variant that leads to misdiagnosis in a clinical context is difficult to quantify. Yet practically this will easily dwarf both the cost of computation and sequence data generation.

Having identified this source of error in variant identification, there may be better ways found to produce reference datasets of RFP variation that can routinely be filtered from variant call sets. As we have shown, these will be specific to the ethnic background of any given genome and argues for the ascertainment of reference population datasets of genomes from diverse ethnic groups. While any reference dataset of RFP variants for a given ethnic group, unless exhaustively ascertained, will almost certainly be incomplete. However, it is evident from our analysis that the sequences of other members of a population, even if only a handful, will likely catalogue the majority of the most prevalent RFP variants specific to a given population. This by itself, without multiple rounds of resampling, realignment and recalling on a given genome, may be sufficient to ameliorate the risk of RFP variant-related misdiagnosis to acceptable levels. Generation of such RFP variant reference sets might be most efficiently performed by databases of genomic variation on a routine basis, and comparison of a given personal genome to this data corpus will annotate variation likely to be due to read misalignment.

## Conclusions

Sequence variation identified by short-read resequencing includes recurrent false positive miscalls, which arise due to read misalignment to redundant regions with high sequence identity. Variants called due to read misalignment are recurrent and can be catalogued for any given genome sequence. Miscalled variants catalogued for diverse individual sequences are show to be almost entirely present in growing population databases of human genomic information. The recurrent false positive variants miscalled in any given genome can be removed by two non-exclusive strategies: i) through cataloguing intermittent variant calls for a single genome or exome, by a resampling, realignment and recalling procedure, and ii) by excluding calls that are not replicated by multiple variant calling tools, in this case, both GATK and SAMtools.

## List of Abbreviations

SNV –: single nucleotide variant
RFP –: recurrent false positive
ENU –: N-ethyl-N-nitrosourea

## Declarations

### Ethics approval and consent to participate

Genome data from Omani individuals were obtained with hospital-based consent to N Al Sukaiti (Royal Hospital, Muscat) for the genetic analysis of these individuals. Indigenous Australian genome data was obtained with appropriate consent granted to SJ Foote (The Australian National University 2014/663). Data from public genome sequences (NA12878) was obtained from the 1000 Genomes file server (ftp://ftp.1000genomes.ebi.ac.uk).

Mouse exome data was obtained under ethics approval 2014/61 (Production and phenotyping of exome sequences ENU gene variant mouse pedigrees) to Dr E Bertram of the Australian National University.

### Consent for publication

Not applicable

### Availability of data and material

An aggregate dataset of variants identified in ENU mutagenized is available from the Australian Phenomics Facility (https://databases.apf.edu.au/download/) and includes the possibility of reanimating cryopreserved strains of interest. Exome sequences for particular mouse strains are available on request to TDA.

Human data analysed in the study is not publically available and controlled under the relevant ethics approvals covering this data, in rare circumstances the data may available from the corresponding author and approval holders, given sufficient justification.

### Authors' contributions

TDA, MAF and GB designed the study, TDA and MAF performed the analysis, MCC, NAS and SJF provided data and TDA and MAF wrote the manuscript.

## Acknowledgements

The authors thank CC Goodnow and the Australian Phenomics Facility for ongoing access to mouse exome datasets collected over many years. We thank the National Computational Infrastructure (Australia) for continued access to significant computation resources and technical expertise.

